# Probabilistic Assessment of Nerve Regeneration with Diffusion MRI: Validation in Rat Models of Peripheral Nerve Trauma

**DOI:** 10.1101/707646

**Authors:** Isaac V. Manzanera Esteve, Angel F. Farinas, Alonda C. Pollins, Marlieke E. Nussenbaum, Nancy L. Cardwell, Hakmook Kang, Mark D. Does, Wesley P. Thayer, Richard D. Dortch

## Abstract

Nerve regeneration after injury must occur in a timely fashion to restore function. Unfortunately, current methods (e.g., electrophysiology) provide limited information following trauma, resulting in delayed management and suboptimal outcomes. Herein, we evaluated the ability of diffusion MRI to monitor nerve regeneration after injury/repair. Sprague-Dawley rats were divided into three treatment groups (sham=21, crush=23, cut/repair=19) and *ex vivo* diffusion tensor imaging (DTI) and diffusion kurtosis imaging (DKI) was performed 1-12 weeks post-surgery. Behavioral data showed a distinction between crush and cut/repair nerves at 4 weeks. This was consistent with DTI, which found that thresholds based on the ratio of radial and axial diffusivities (RD/AD=0.40±0.02) and fractional anisotropy (FA=0.53±0.01) differentiated crush from cut/repair injuries. By the 12^th^ week, cut/repair nerves whose behavioral data indicated a partial recovery were below the RD/AD threshold (and above the FA threshold), while nerves that did not recover were on the opposite side of each threshold. Additional morphometric analysis indicated that DTI-derived normalized scalar indices report on axon density (RD/AD: r=−0.54, p<1e-3; FA: r=0.56, p<1e-3). Interestingly, higher-order DKI analyses did not improve our ability classify recovery. These findings suggest that DTI can distinguish successful/unsuccessful nerve repairs and potentially identify cases that require reoperation.

## INTRODUCTION

Traumatic peripheral nerve injuries (TPNIs) are commonly caused by penetrating injuries, crush mechanisms, stretch, lacerations, and/or ischemia ^1^. Primary repair is recommended when nerve ends are severed without tension ^2–4^; however, approximately 40% of these surgeries fail and require a secondary surgical procedure to restore sensory and motor function ^5^. After repair, nerves regeneration occurs at approximately 1 mm/day, which can result in a lengthy recovery when the injury site is not in close proximity to the neuromuscular junction ^6^. Although electromyography and nerve conduction studies (NCS) are the current standards to assess nerve regeneration in extremities ^7^, these methods are incapable of monitoring and quantifying nerve recovery until muscles are reinnervated. Thus, with no insight into the recovery process, physicians are forced to delay second interventions until the time at which axons should be reaching the muscular end plate ^8^, which increases the probability of irreversible muscular atrophy ^9,10^. Given these limitations, new methods are needed to provide an accurate evaluation of nerve regeneration throughout the recovery process to improve clinical decision-making and outcomes in patients.

Diffusion-weighted magnetic resonance imaging (MRI) probes tissue features at the microstructural level by measuring the effect of tissue barriers on the apparent diffusion of water molecules ^11^. In nerves, the ordered arrangement of axons results in an apparent diffusion coefficient (ADC) that is lower perpendicular to axons than parallel to them ^12–14^. Diffusion tensor imaging (DTI) is a commonly used approach that measures diffusion along multiple directions to quantify indices that describe this diffusion anisotropy, including the mean diffusivity (MD, mean value across all directions), axial diffusivity (AD, diffusivity along axons), radial diffusivity (RD, diffusivity across axons), and fractional anisotropy (FA = 0-1, higher values indicate higher anisotropy). Higher-order diffusion kurtosis imaging (DKI) methods ^15–20^ further quantify the effect of restricted diffusion within axons and/or heterogeneity (e.g., different intra/extra-axonal diffusivities), yielding measures of mean (MK, mean kurtosis across all directions), axial (AK, kurtosis along axons), and radial kurtosis (RK, kurtosis across axons) that are potentially more specific to axon microstructure ^21–24^. For these kurtosis measures, larger values are indicative of increased non-Gaussian diffusion from tissue heterogeneity and/or restricted diffusion.

Based upon its sensitivity to nerve microstructure, diffusion MRI methods have been used to monitor nerve degeneration and/or regeneration in animal models ^25–29^. From these studies, it has been established that DTI metrics longitudinally track with electrodiagnostic and functional assessments of recovery. In addition, RD and FA values have been shown to correlate with behavior and histological measures of axon density during regeneration ^27^. We recently demonstrated that DTI indices also report on injury severity acutely after transection and surgical repair ^8^ and these indices track with behavioral recovery over time ^30^. DKI indices may offer improved specificity to changes in nerve microstructure; however, the relationship between DKI parameters and peripheral nerve regeneration has yet to be validated. Nevertheless, recent work has indicated that nerve DKI is both feasible and sensitive to nerve injury ^31^.

While previous MRI studies have demonstrated the relationship between diffusion MRI indices in nerves and function or histology ^8,25,27,28^, no studies have evaluated the ability of diffusion MRI to stratify successful from unsuccessful repairs after TPNI. In this study, we propose a probabilistic model of nerve recovery based on multiple diffusion MRI parameters. To develop this model, we performed high-resolution *ex-vivo* DTI and DKI at various time-points after *i*) injuries that are self-resolving (crush) and *ii*) injuries that show variable recovery (neurotmesis followed by surgical repair, i.e. cut/repair). Additional sham surgeries were performed for comparison. From these data, a probabilistic model of nerve recovery was developed and validated against behavioral and pathological findings. In addition, cut-off values that distinguish successful/unsuccessful repairs were determined, with the long-term goal of using these cut-off values to identify cases that require reoperation.

## RESULTS

### Diffusion and Kurtosis Parameter Maps

Figure 1 shows representative images and parameter maps from a distal slice in cut/repair nerves 12 weeks after recovered (top panel) and non-recovered nerves (bottom panel) based on behavioral findings. Note the elevated diffusivities (AD, RD, and MD), reduced kurtosis indices (AK, RK, and MK), and reduced FA values in the non-recovered nerve relative to the recovered nerve. In addition, note the heterogeneity in parameter maps of the non-recovered nerve relative to the recovered nerve, which may be indicative of partial regeneration in this nerve.

**Figure 1.**
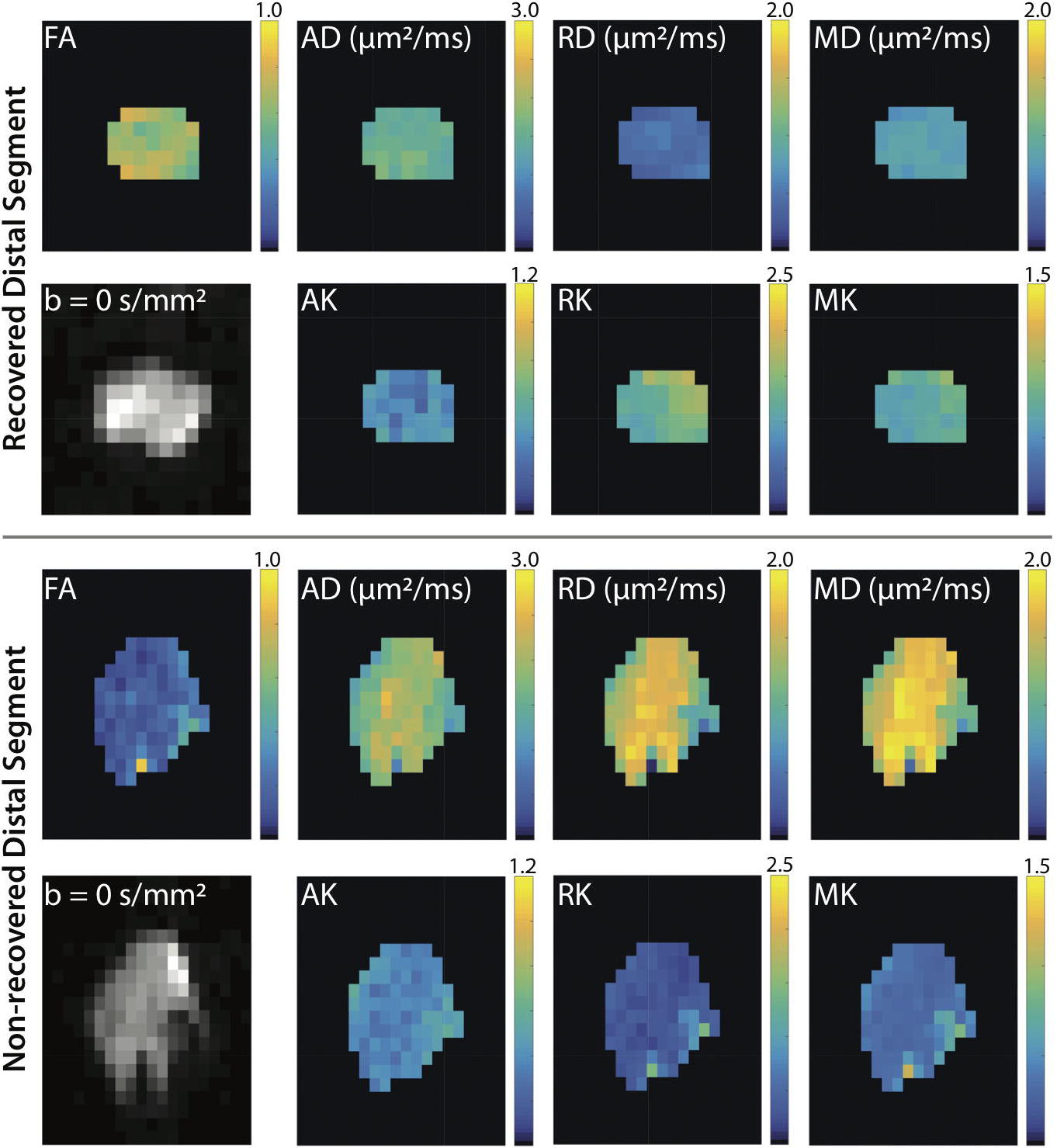
Representative distal DTI/DKI parameter maps from recovered (top panel) and non-recovered (bottom panel) cut/repair nerves 12 weeks after injury. Shown are maps of DTI (FA, AD, RD, and MD) and DKI (AK, RK, and MK) parameters along the non-diffusion weighted image (b=0 s/mm^2^) from a single distal slice. Note the reduced FA (blue hues) and elevated diffusivities (yellow hues) in the non-recovered nerve relative the recovered nerve. In addition, note the reduce kurtosis measures (blue hues) in the non-recovered nerve.

### Evolution of Diffusion and Kurtosis Parameters Following Surgery

Figure 2 summarizes the evolution of the mean distal diffusion/kurtosis parameter for each cohort after surgery. Changes in diffusion parameters in the early and late timepoints can be linked to inflammatory/degenerative and regenerative processes, respectively ^8,11,13,20,26^. With this in mind, RD values decreased in a similar fashion after surgery in crush and sham nerves, while a slower and less substantial decrease in RD was observed in cut/repair nerves (p<1e-3 for 2, 4, and 12 weeks), which may be due to heterogeneity within in this cohort (i.e., not all cut/repair nerves recovered). In contrast, AD differed across all three cohorts. In particular, a significant decrease in AD was observed at two weeks after crush injury relative to both sham (p<1e-3) and cut/repair cohorts (p<1e-3), suggesting that AD and RD are sensitive to different pathological features during the recovery process. Because MD was formed from a weighted average of AD and RD, it did not provide unique information. The kurtosis parameters exhibited an inverse trend relative to each DTI parameter, although the temporal evolution may differ between diffusion and kurtosis parameters (e.g., RD reached a maximum at two weeks, while RK reached a minimum at four weeks in cut/repair nerves).

**Figure 2.**
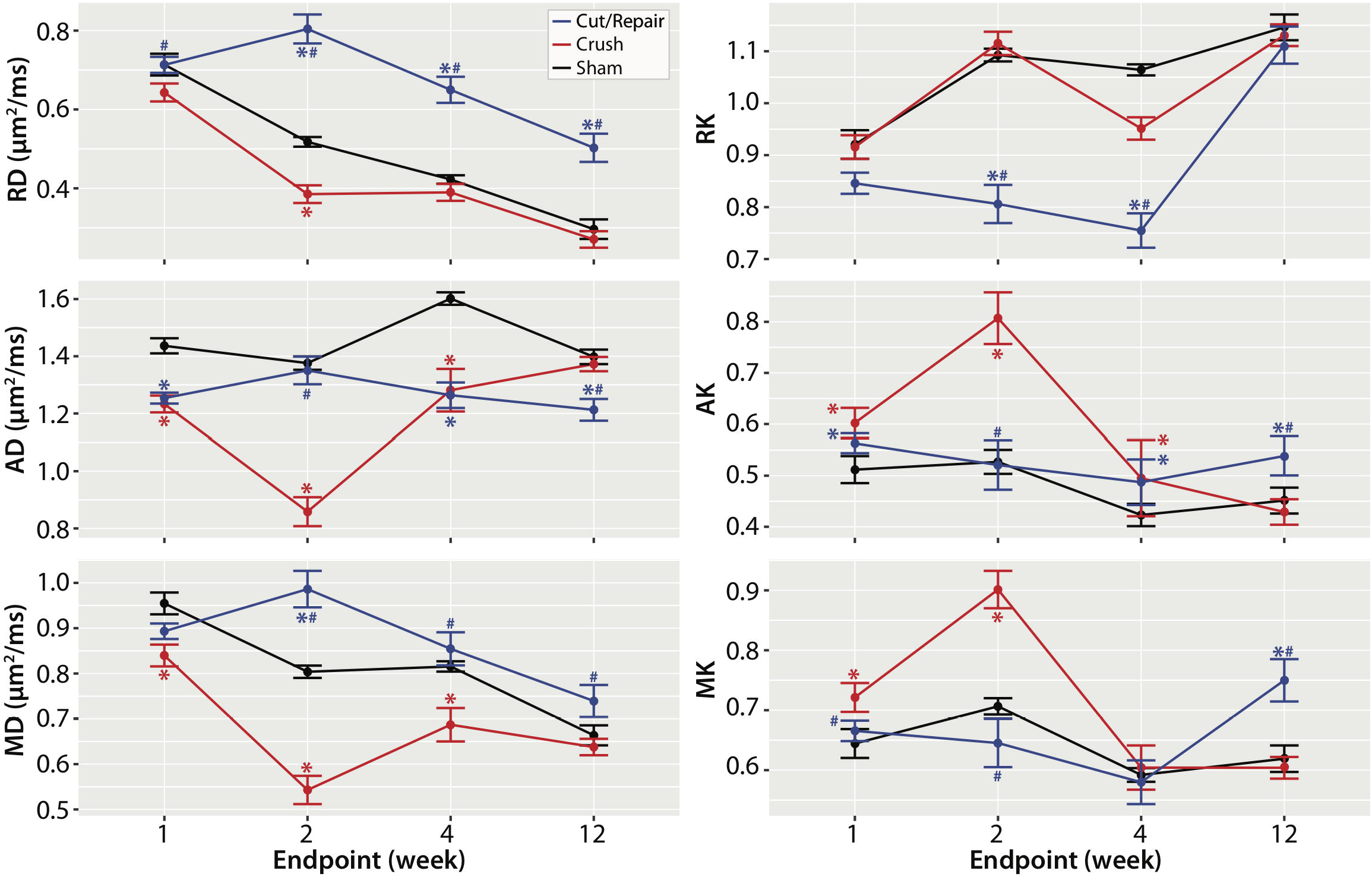
Evolution of diffusivities (left column) and kurtosis parameters (right column) after each injury/surgical intervention. Changes in parameters at 1 and 2 weeks are influenced by inflammation/edema, while changes at weeks 4 and 12 are indicative of de/regeneration. RD values decreased in a similar fashion after surgery in crush and sham nerves, while a slower and less substantial decrease in RD was observed in cut/repair nerves. In contrast, AD differed across all three cohorts, but were similar in crush and cut/repair nerves after four weeks. Finally, the kurtosis parameters exhibited a similar, but inverse, trend relative to each diffusion parameter. Significant differences at each time-point are indicated as follows: cut/repair vs. sham = blue asterisk, crush vs. sham = red asterisk, cut/repair vs. crush = blue hashtag.

### Probabilistic Model of Recovery Based on Diffusion and Kurtosis Parameters

In developing a probabilistic model of nerve recovery, we analyzed the ability of combinations of DTI and DKI parameters to differentiate between self-resolving crush injuries and partially recovering cut/repair nerve injuries. Of particular note, we analyzed slice-wise radial vs. axial parameters as shown in Figure 3. During the first two weeks, neither DTI nor DKI parameters demonstrated a clear differentiation between injury types, likely due to the dominant effect of edema. By the fourth week however, RD vs. AD plots yielded non-overlapping clusters for each cohort. Note that this separation was not seen in the counterpart RK vs. AK plot; therefore, we limit our discussion below to RD vs. AD. The contrast observed in the RD vs. AD plot at four weeks is consistent with the longitudinal behavioral data in Figure 4, which indicated that the crush nerves were fully recovered by four weeks while cut/repair nerves were not. Based upon this differentiation and the assumption that regeneration is complete 12 weeks after surgery, we predicted that the cut/repair nerves whose RD/AD values overlap with crush nerves at 12 were succesful repairs, while those that did not overlap with were unsuccesful repairs.

**Figure 3.**
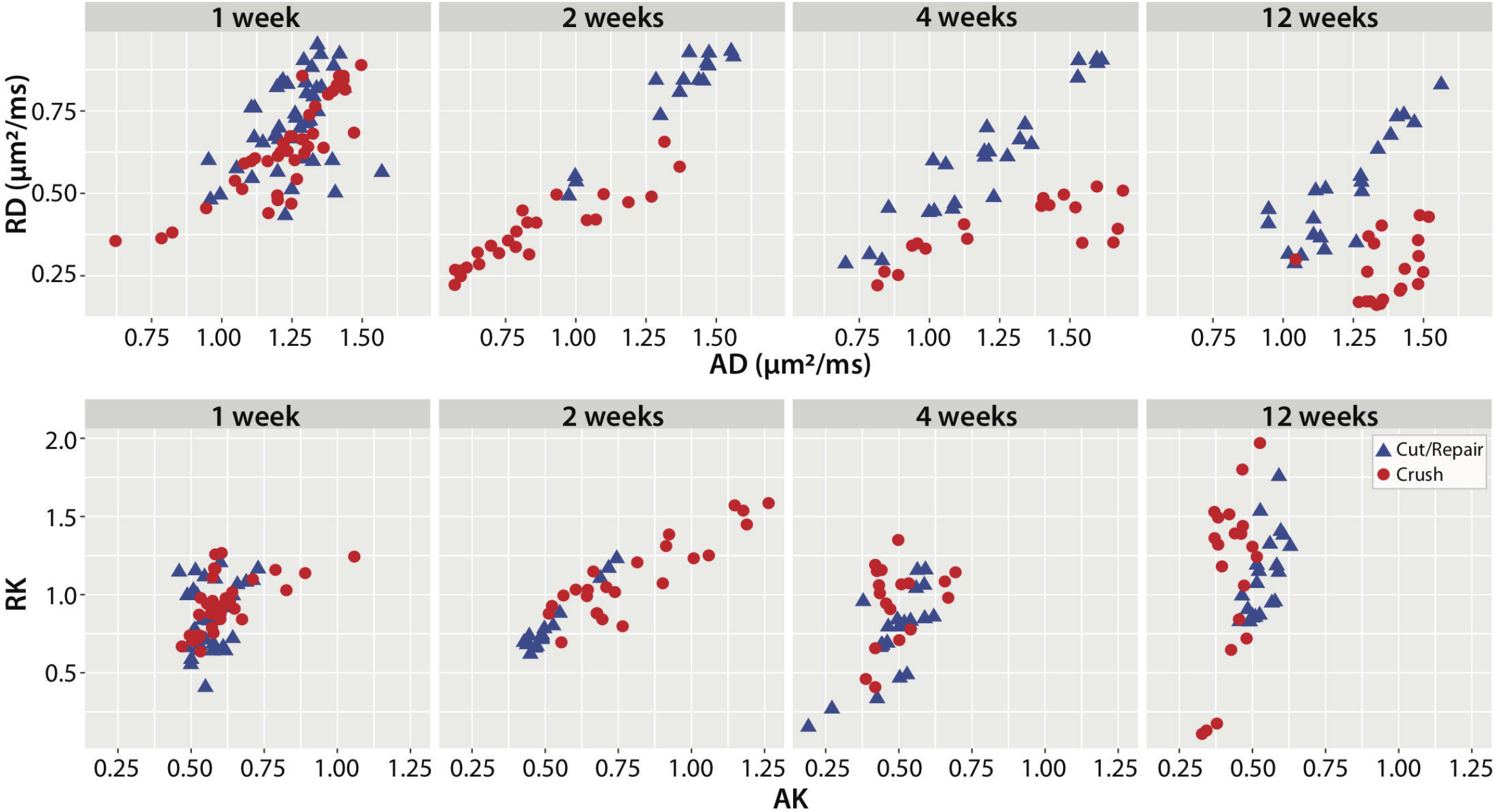
Scatterplots of distal radial vs. axial diffusivities (top panel) and kurtosis indices (bottom panel) for crush and cut/repair nerves at each time point after injury (left-to-right). Each point represents the mean cross-sectional value for a slice distal to the injury site.

**Figure 4.**
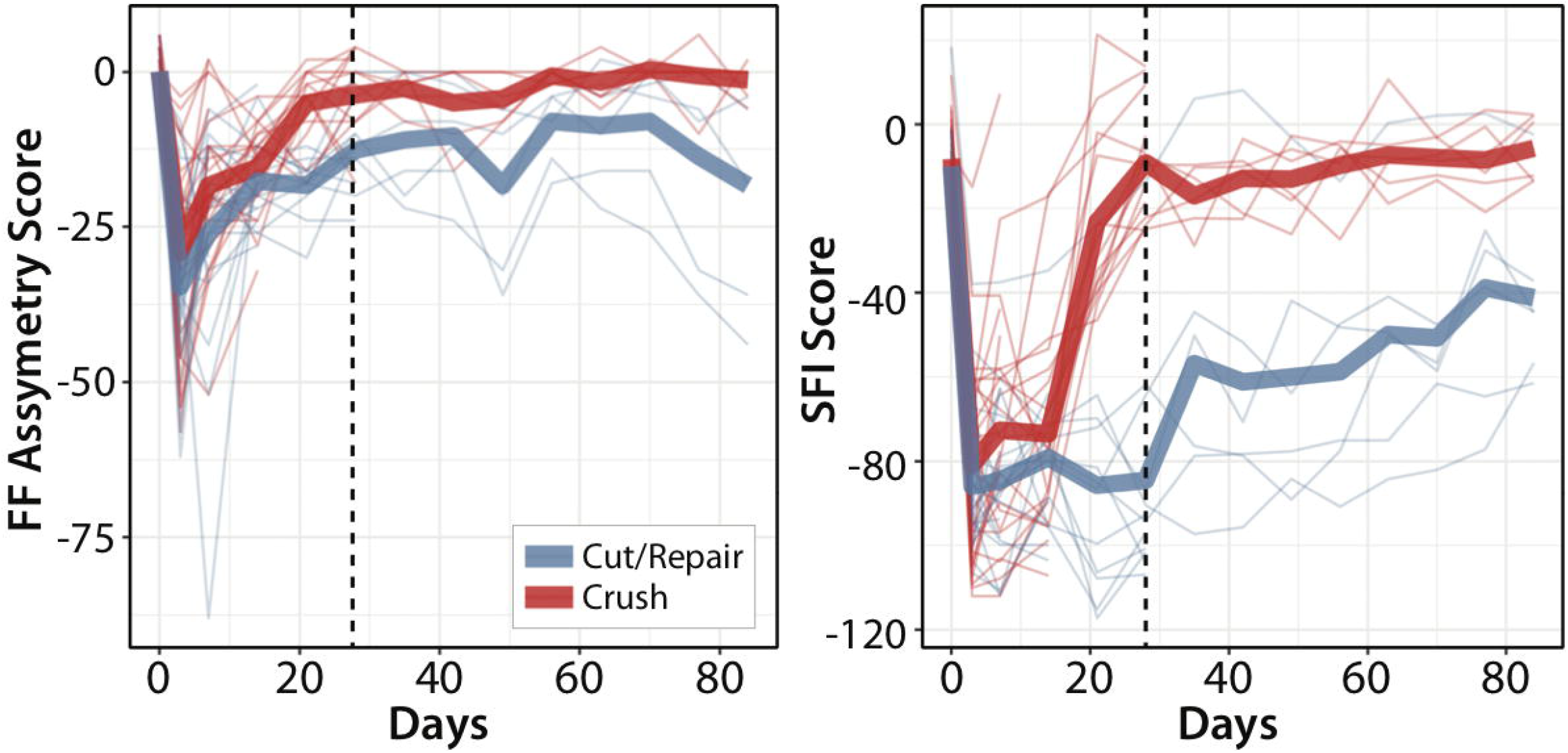
Longitudinal behavioral data after crush (red lines) and cut/repair injuries (blue lines). Results are shown for both FF (left panel) and SFI (right panel). The longitudinal time course (from injury to euthanasia) for each individual animal is plotted with thin lines, while the mean time course within each cohort is plotted with thick lines. Note the sharp decay in FF and SFI for both cohorts immediately after surgery, with the largest contrast between injury types at 4 weeks (vertical black dashed line). These results indicate a faster behavioral recovery following crush relative to cut/repair injuries, with crush injuries nearly recovered by four weeks.

To generate this predictor, the linear boundary between crush and cut/repair nerves was calculated for distal RD vs. AD values at four weeks using SVM analysis (see *Methods*). The decision boundary (RD/AD=0.40±0.02) separated recovered nerves (crush) from non-recovered nerves (transected/repair) four weeks after injury. As expected, the intercept of this decision boundary was close to zero (− 0.03±0.01).

This decision boundary was then tested to the other distal time-points (1, 2, and 12 weeks) as well as proximal nerve segments and sham nerves for all time-points (1, 2, 4, and 12 weeks). Scatterplots of the results color-coded by probability of recovery (see *Methods*) can be seen in Figure 5. Once again, distal segments of injured nerves (top row) were dominated by inflammation/edema in the first two weeks, resulting in low recovery probabilities. By the fourth week, behavioral data (Figure 4) indicated that crush injuries were fully recovered, while cut/repair injuries were not yet recovered. This is reflected in Figure 5 where recovery probabilities were above 50% for all crush injuries and below 50% for all cut/repair injuries. At 12 weeks, however, a subset of cut/repair nerves had a high probability of recovery, while other nerves had a low probability of recovery (see below for additional analysis of these individual samples). Results from the proximal segments and sham nerves (bottom row) further validated this model, with increasing recovery probabilities as edema subsided and nearly full recovery by 4 and 12 weeks.

**Figure 5.**
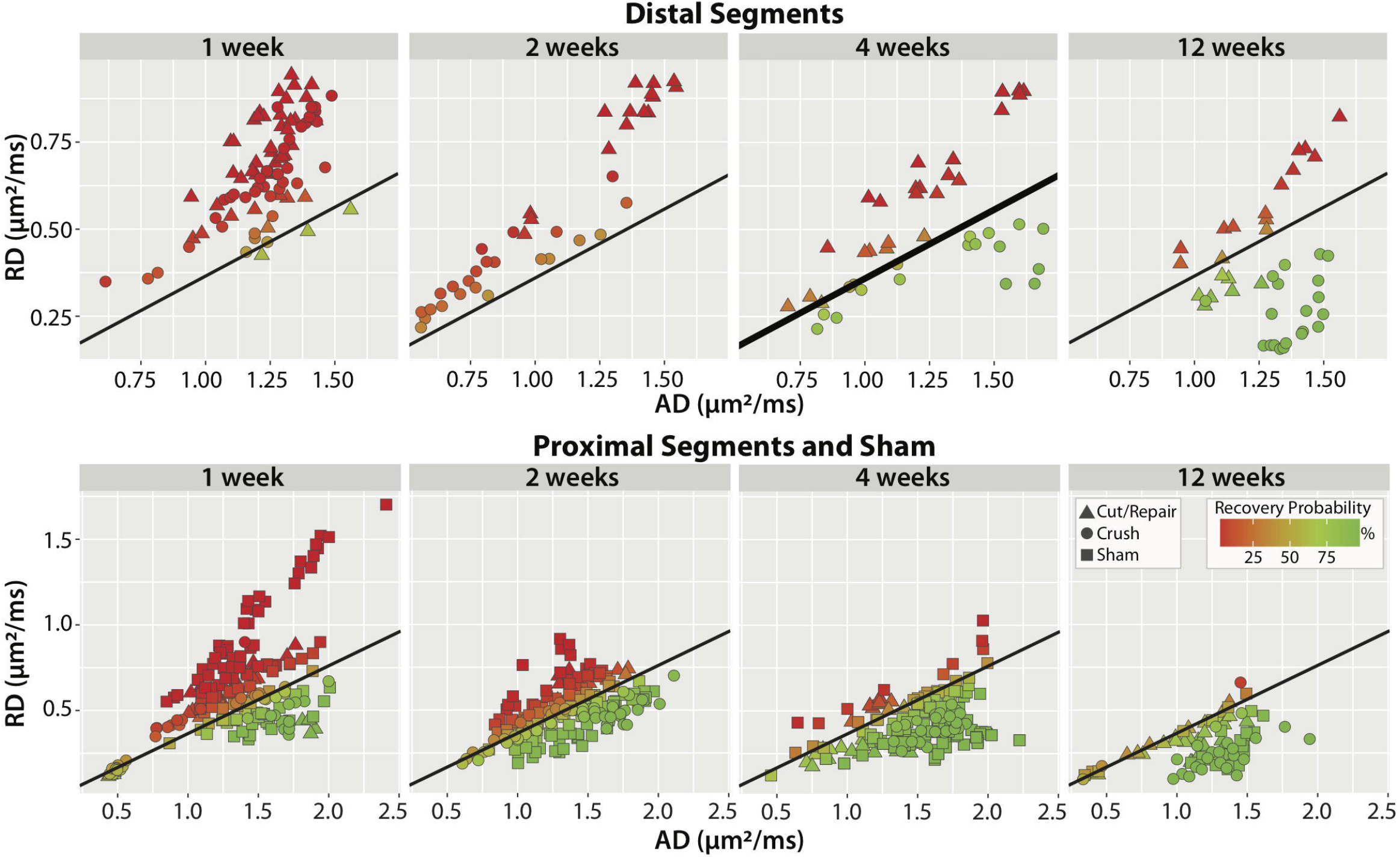
Scatterplots of radial vs. axial diffusivities for distal segments (top panel), proximal segments (bottom panel), and sham nerves (bottom panel). The SVM boundary was generated using the distal data at four weeks (thick black line) and tested using data from the other panels (thin black line). Each point represents the mean cross-sectional (or slice-wise) value. Recovery probabilities are color-coded from non-recovery (red, above the SVM boundary) to recovery (green, below the SVM boundary). Consistent with the behavioral findings in Figure 4, the model indicates that crush injuries are recovered distally by the 4^th^ week, while cut/repair nerves are not. By the 12^th^ week, however, cut/repair nerves are found on both sides of the decision boundary (green/red triangles), indicating different probabilities of recovery. This model was further tested in sham nerves and the proximal regions of injured nerves (bottom panel), nearly all of which showed a high probability of recovery by the 4^th^ week.

### Converting RD/AD to an FA Threshold

Although the ratio RD/AD stratifies crush and cut/repair nerves at 4 weeks, it is not a conventionally reported DTI metric. As a result, we transformed RD/AD to a corresponding FA cut-off using the following relationship ^32^

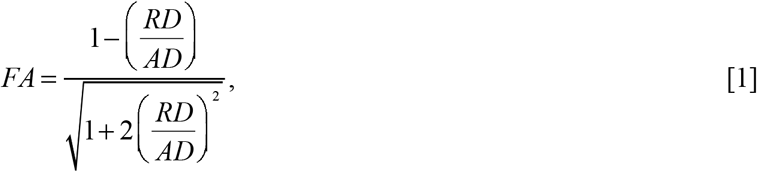

which assumes a cylindrically symmetric tensor. This indicates that a linear RD/AD boundary with zero offset can be represented by a unique FA “success threshold” (FAST)=0.53±0.01. To test this FAST value, Figure 6 shows distal slice-wise FA profiles 12 weeks after injury for six samples stratified by behavioral recovery: two fully recovered crush nerves, two non-recovered cut/repair nerves, and two fully recovered cut nerves. It can be seen that FA values along the length of both crush nerves are considerably above the FAST value, in agreement with the statement that all crush injuries fully recover. FA values for the distal regions of cut/repair nerve injuries that recovered were also above the FAST value, while nerves with failed recoveries display values below this threshold. In addition, note the reduction in FA near the injury site for the crush and cut/repair nerves even 12 weeks after injury. This may be due to nerve sprouting and a reduction in coherence of axons that takes place at the injury site ^10,28,33^. In addition, since nerve growth is not homogeneous along the nerve, we can expect the density of fibers to decrease distal to the injury, which may be reflected by the reduced FA values in the most distal regions of the recovered cut/repair nerves.

**Figure 6.**
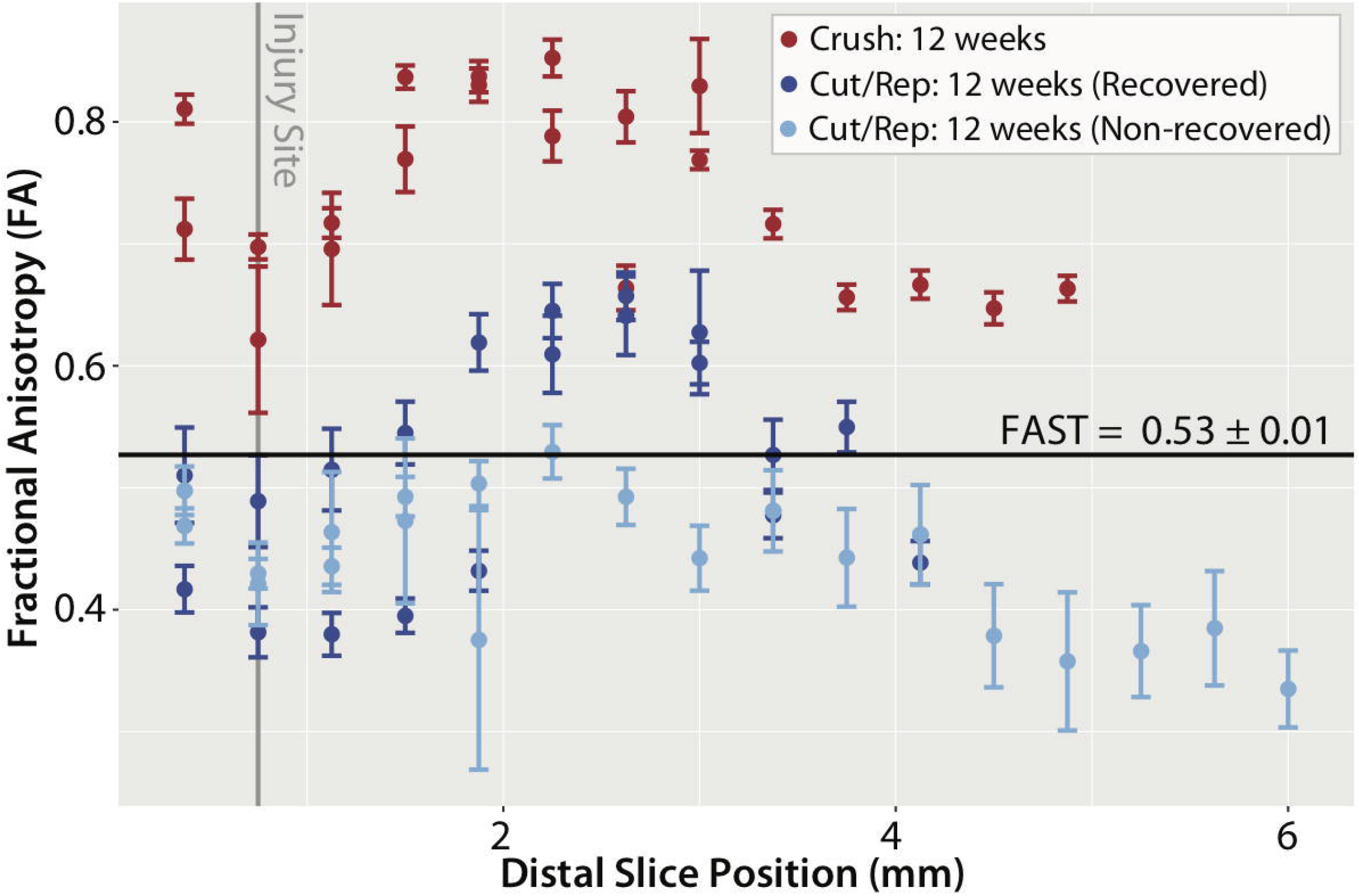
Slice-wise FA values (mean ± SEM) distal to the injury site (vertical gray line) for six nerve samples at 12 weeks after each intervention. Shown are representative results from two crush samples (red) along with the four cut/repair samples, two of which showed evidence of recovery via behavioral measures (dark blue) and two of which did not show evidence of recovery (light blue). In the recovered nerves, mean FA values were above the FAST line for a majority of distal slices. In contrast, mean FA values were below this cut-off value for almost all slices in the non-recovered nerves.

### Comparison on DTI/DKI and Histology

Figure 7 shows representative histological images at 2, 4 and 12 weeks for the three cohorts. Correlations between DTI/DKI parameters in distal nerves and the resulting histologically-derived measures of axon density are shown in Figure 8. Note the significant correlation between radial indices (RD: r=−0.49, p<1e-3 and RK: r=0.40, p<1e-2) and axon density. In contrast, no correlation was overserved between the axial indices (AD: r=0.10, p=0.49 and AK: r=−0.22, p=0.14) and axon density. Consistent with our MRI findings in Figure 4, DTI-derived normalized scalar indices (RD/AD: r=−0.54, p<1e-3 and FA: r=056, p<1e-3 exhibited the strongest relationship with axon density.

**Figure 7.**
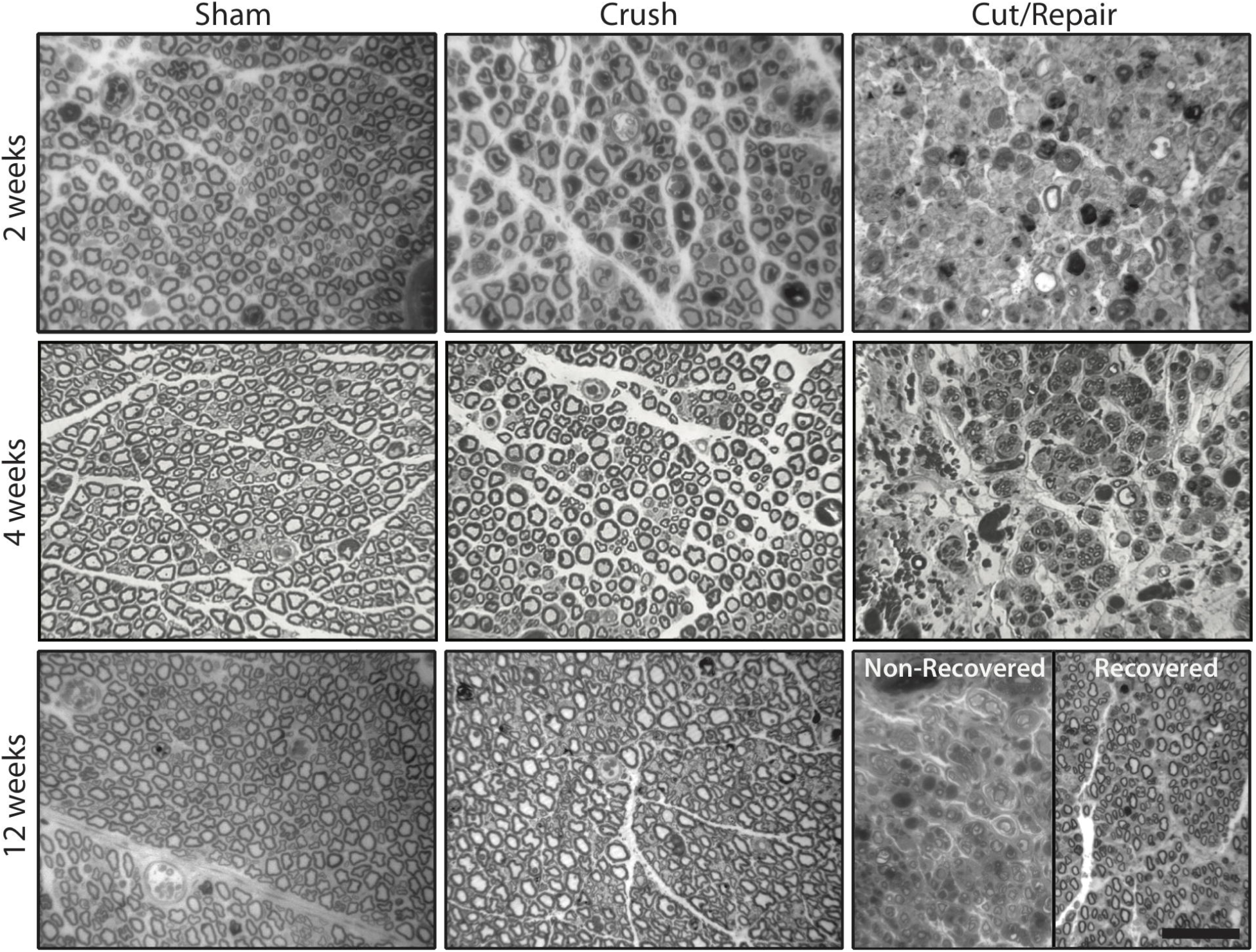
Representative distal toluidine blue stained sections at 2 (top panel), 4 (middle panel) and 12 weeks (bottom panel) after each injury type. In the sham groups (left column), there was little difference between the three times, except for some evidence of extra-cellular edema at 2 weeks. In contrast, crush nerves (middle column) shows indications of degeneration at 2 weeks and full recovery by 4 weeks. For the cut/repair group (right column), severe degeneration is observed at 2 and 4 weeks. By 12 weeks, a subset of cut/repair nerves showed little recovery (“non-recovered” panel), while another subset showed partial recovery (“recovered” panel). The scale bar in the bottom-right panel represents 50 μm.

**Figure 8.**
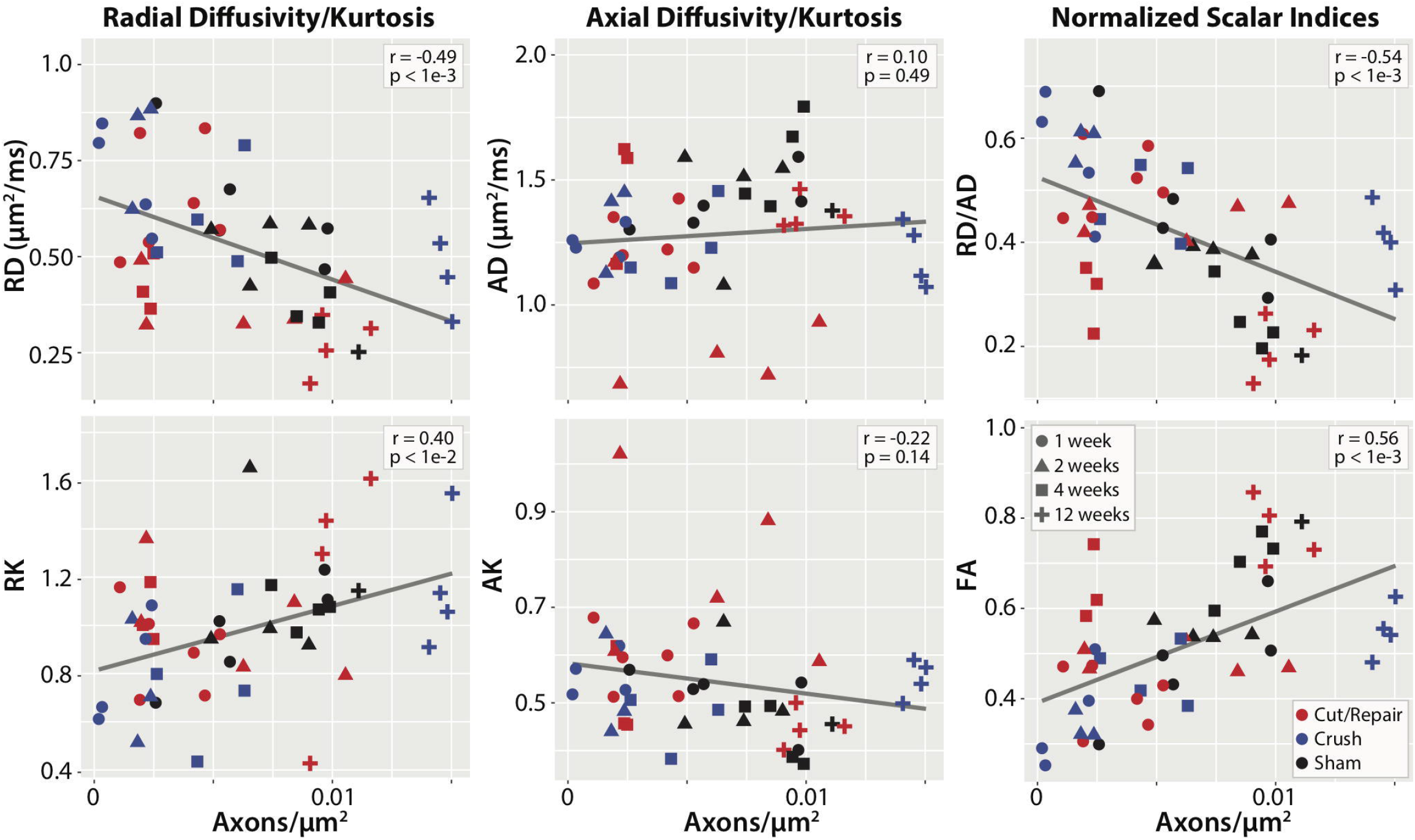
Linear correlations between DTI/DKI parameters and histologically-estimated cross-sectional axon densities. Results are shown for all time-points (encoded via marker shape) and injury type (encoded via marker color). Significant correlations were observed between radial parameters (RD and RK) and axon density (left column), while the relationship between axial parameters (AD and AK) and axon density (middle column) was not significant. The normalized scalar indices from DTI (right column, RD/AD and FA) demonstrated the strongest relationships with axon density. The Pearson’s correlation coefficient (*r*) and corresponding *p*-value is given in the white box of each panel.

## DISCUSSION

We demonstrated that diffusion MRI can successfully identify recovered from non-recovered nerves 4 weeks after cut injuries and surgical repair. Consistent with previous studies ^27^, behavioral findings (Fig. 4) indicated that the crush nerves were fully recovered approximately 4 weeks after injury, while the cut/repair nerves did not substantially recover until after four weeks. In other words, a clear distinction between the behavioral recovery of crush and cut/repair nerves was observed at 4 weeks, which is consistent with the RD/AD measurements at the same timepoint in Fig. 3. Based upon this information, we developed a statistical model of recovery using distal RD/AD values 4 weeks after injury and surgical repair. As shown in Fig. 5, the linear model successfully classified the crush and cut/repair nerves at 4 weeks. By the 12^th^ week, cut/repair nerves were found on both sides of the classification boundary, indicating different likelihoods of recovery. When further tested in sham nerves and the proximal regions of injured nerves, nearly all regions showed a high probability of recovery by the 4^th^ week as expected. RD/AD values were then converted to more commonly reported FA values and an FA “success threshold”, or FAST, was tabulated using Eq. [1]. As shown in Fig. 5, the cut/repair nerves that recovered (based on behavioral findings) had slice-wise FA values were that were above the FAST line for a majority of distal slices. In contrast, FA values were below this cut-off value for almost all distal slices in the non-recovered nerves. To validate this model histologically, light microscopy was performed in the distal regions of these same nerves and cross-sectional axon densities were manually estimated (Figs. 7–8). Consistent with previous work in crush injured nerves ^27^, both normalized scalar indices (RD/AD and FA) reported on changes in axon density during the recovery process for all three cohorts (sham, crush, and cut/repair).

In addition to nerve de/regeneration, the normalized scalar indices were influenced by edema within the first two weeks, which can be visualized as increased extra-cellular space in the light micrographs (Fig. 7). Because the model was trained using data after edema subsided (4 weeks), it incorrectly predicted that all nerves following crush injury had a low probability of recovery at 1-2 weeks (Fig. 5). This limitation was further verified using data from sham nerves and the proximal regions of crush nerves at 1-2 weeks, all of which were incorrectly classified as low probability recoveries. Given this limitation, alternate models based on other diffusion parameters may be necessary to classify nerve recovery early after injury. For example, previous work ^34–36^ has shown that axial diffusion parameters are primarily sensitive to axonal degeneration within the first few weeks after injury, which is consistent with the observed decrease in AD (and increase in AK) in crush nerves at 2 weeks (Fig. 2). In addition, diffusion models that account for the effects of edema ^12^ or filter out signal contributions from the extra-cellular space ^37^ may improve predictions of nerve recovery within the first few weeks.

One interesting finding of this study was that conventional DTI, without the need for a higher-order DKI model, was able to differentiate recovered from non-recovered nerves following injury and repair. Previous work in the central nervous system ^21–24^ suggested that DKI indices may provide a more specific assay of axon microstructure than DTI, which is in conflict with our findings in peripheral nerves. These differences may be attributed in part to the different b-values (2000 and 4000 s/mm^2^) used herein, which were empirically determined herein to account for the decreased diffusivity caused by fixation ^38^ and temperature (bore temperature ≈ 20°C). Perhaps more significant, axon diameters in peripheral nerves (≈1-15 μm)^39^ are nearly an order of magnitude larger than in the central nervous system, suggesting that longer diffusion times (Δ=12 ms for the current study) may be required for optimal sensitivity to diffusion kurtosis in the presence of these larger axons ^40^. For example, axons are often modeled using the zero-radius approximation, which assumes Δ ≫ *R*^2^/*D*_a_, where *R* is the radius of the axon and *D*_a_≈1 μm^2^/ms is the free intra-axonal diffusivity ^41^. For white matter data acquired on a clinical system (Δ≈50 ms, *R*≈1 μm), the zero radius-approximation is valid. In contrast, this assumption is no longer valid for peripheral nerve data acquired herein (Δ=12 ms, *R*≈5 μm). Given these differences, additional studies on the relative effects of restricted diffusion within axons and heterogeneity (i.e., differences in intra/extra-axonal diffusivities) on DKI parameters in nerves are warranted. Nevertheless, the ability to predict recovery via DTI is beneficial for clinical translation, as DKI requires longer scans (i.e., additional b-values and/or directions) than conventional DTI.

Although promising, two additional limitations of this study should be addressed in future work. First, *ex vivo* MRI was performed in fixed rather than fresh tissues due to the long scan times required for high-resolution MRI. Because fixation affects diffusivities, one cannot directly translate the absolute diffusivities reported herein to fresh tissues. Fortunately, previous studies ^38^ have indicated that normalized scalar indices (RD/AD and FA) are equivalent in fresh and fixed tissues. This suggests that the model proposed herein may be applicable *in vivo*, although additional studies are required to confirm this. Second, the current study investigated a cross-section of samples at several timepoints (1, 2, 4, and 12). Future studies should focus on *i*) additional timepoints between 4-12 weeks to determine how early our model can stratify recovered from non-recovered nerves and *ii*) longitudinal *in vivo* measurements to determine how model predictions relate to outcomes in the same nerve.

In conclusion, normalized scalar indices (RD/AD and FA) derived from high-resolution *ex vivo* DTI of rat sciatic nerve were found to differentiate crush (fully recovered) from cut/repair (partially recovered) injuries once the edema had subsided at 4 weeks post-injury. In addition, these indices related to behavioral recovery and histological measurements of axon density. Interestingly, higher order DKI models, which require longer scan times than DTI, did not improve the sensitivity of the model to regeneration. Currently, nerve recovery after surgical intervention requires a “wait and watch” approach due to the absence of a reliable noninvasive biomarker, which increases the likelihood or poor outcomes. The studies herein suggest that RD/AD and FA can distinguish successful/unsuccessful repairs earlier than currently possible and potentially identify cases that require reoperation, although additional *in vivo* longitudinal studies are required to determine how model predictions relate to long-term outcomes.

## METHODS

### Ethics Approval

All animal procedures were approved by the Vanderbilt University Medical Center Institutional Animal Care and Use Committee, under the Guide for Care and Use of Laboratory Animals to minimize pain and suffering.

### Animal Grouping

Sixty-three Sprague-Dawley rats were used as experimental animals. The rats were divided into groups accordingly to their traumatic peripheral nerve injury. Animals were euthanized at different time points after each intervention (1, 2, 4, and 12 weeks) to perform ex-vivo MRI scanning.

### Behavioral Testing

Behavioral tests were performed before surgical intervention, three days after intervention, and then weekly until each animal’s endpoint. Measurements included foot fault (FF) asymmetry score and sciatic function index (SFI), as described in previous studies ^42^.

### Experimental Treatment

Induction and general anesthesia were performed with a dose of 3 mL/min of 2% Isoflurane, and care was taken the potential for hypothermia. A 3-cm skin incision was then made from below the ischial notch parallel the longitudinal axis of the hind leg. Dissection was done by planes to free the sciatic nerve proximally and distally up to its trifurcation. Within this nerve segment and 1cm proximal to the trifurcation, each animal received one of the following: application of a Hemostat for 10 seconds (crush, *n* = 23), full transection with immediate repair (cut/repair, *n* = 19), or no intervention (sham, *n* = 21). Repair surgery consisted of an end-to-end fashion with interrupted epineurial 9-0 nylon sutures (Ethicon, Somerville, NJ). Wounds were closed in 2 layers using 5-0 Monocryl suture (Ethicon, Somerville, NJ). After surgery, animals were carefully monitored for any adverse anesthetic effect and provided with daily injections of ketoprofen (5mg/kg) for 3 days post-operatively. At each animal’s endpoint, euthanasia was performed with an intracardiac dose of 120 mg/kg of Euthasol (Virbac AH, Fort Worth, Texas), and nerves were harvested.

### Tissue Sample Preparation

Harvested nerves were fixed by immersion in 3% glutaraldehyde/2% paraformaldehyde, washed for a minimum of 1 week in phosphate-buffered saline (PBS) to remove excess fixative, then immersed in 1 mM Gd-DTPA (Magnevist; Berlex, Montville, NJ) at 4 °C for at least 36 hours to reduce spin-lattice relaxation times and corresponding scan times. Samples were trimmed to approximately 1 cm in length with crush and cut/repair regions at the center of each segment. Finally, nerves were placed in 1.75-mm glass capillary tubes filled with a perfluorcarbon solution (Fomblin; Solvay, Thorofare, NJ) to prevent tissue drying without contributing to the proton MRI signal.

### MRI Protocol

To improve throughput, groups of six nerves were arranged in a hexagonal pattern and scanned simultaneously. Diffusion-weighted MRI data were acquired at bore temperature (≈20°C) using a 7-T, 16 cm bore Bruker Biospec console (Rheinstetten, Germany) and a 25-mm quadrature radio-frequency coil (Doty Scientific, Columbia, SC) for transmission and reception. Images were acquired with a three-dimensional diffusion-weighted spin-echo sequence and the following parameters: field-of-view=60×60×160 mm^3^, resolution=125×125×372 μm^3^, TE/TR=22/425 ms, gradient pulse duration/diffusion time (δ/Δ)=4/12 ms, b-values=2000 and 4000 s/mm^2^, 20 diffusion directions, number of averaged excitations=2, and scan time = 7 hours and 40 minutes for each b-value.

### MRI Analysis

Diffusion and kurtosis tensors were estimated on voxel-wise basis using a weighted linear least-squares estimation ^19^ in MATLAB 2017b (Mathworks, Natick, MA). The following indices were estimated from the diffusion tensor: FA, MD, RD, AD. From the kurtosis tensor, we additionally estimated MK, AK, and RK. Each slice was manually classified as: proximal to injury, within zone of injury, or distal to injury; and regions-of-interest (ROI) were drawn manually in each nerve to calculate the mean slice-wise diffusion parameters.

### Histology

Consecutively after MRI scans, samples were fixed in 3% glutaraldehyde/2% paraformaldehyde, counter-stained with 1% OsO4 solution, dehydrated in increasing concentrations of ethanol, and embedded in resin. Sections of 1μm were obtained from the center of the distal segment then stained with 1% toluidine blue for examination with light microscopy (Olympus Vanox-T AH-2). Axon counts and cross-sectional areas were manually measured at 40x magnification using Image Pro Plus 7.0 (Media Cybernetics, Bethesda, MD), and axon density was estimated from the number of axons/cross-sectional area. This process was repeated in three randomly selected areas per sample and the mean axon density was reported.

### Statistical Analysis

All statistical analyses were performed R version 3.3.2. For each timepoint and DTI/DKI parameter, pairwise differences between cohorts (sham, crush, cut/repair) were assessed using an unpaired student’s t-test. Correlations between DTI/DKI parameters and histologically-derived axon densities were quantified via Pearson’s correlation coefficient. All p-values were adjusted for the effect of multiple comparisons using a false discovery rate approach ^43^.

A linear classifier for successful versus unsuccessful regenerations was generated for RD/AD values, which showed a clear separation between crush and cut/repair cohorts at 4 weeks after injury/repair. This linear classifier was generated via a support vector machine (SVM library e1071), and uncertainties in the classification boundary were calculated via Bootstrap. This boundary was then tested using distal RD/AD values at the other timepoints (1, 2, and 12 weeks). Additional testing was performed by applying the classification boundary to RD/AD values from sham nerves and the proximal segments of crush and cut/repair nerves at all timepoints (1, 2, 4, and 12 weeks). *A posteriori* classification probabilities were then estimated using an improved implementation of Platt’s approach ^44,45^. This approach was employed rather than a direct classification using logistic regression because the cohorts were fully separable in the training data (distal RD/AD values at 4 weeks), which results in a degenerate solution when logistic regression is employed for classification purposes.

